# Uncertainty modulates visual maps during non-instrumental information demand

**DOI:** 10.1101/2021.07.20.453107

**Authors:** Yvonne Li, Nabil Daddaoua, Mattias Horan, Jacqueline Gottlieb

## Abstract

Animals are intrinsically motivated to resolve uncertainty and predict future events. This motivation is encoded in cortical and subcortical structures, but a key open question is how it generates concrete policies for attending to informative stimuli. We examined this question using neural recordings in the monkey lateral intraparietal area (LIP), a visual area implicated in attention and gaze, during non-instrumental information demand. We show that the uncertainty that was resolved by a visual cue enhanced visuo-spatial responses of LIP cells independently of reward probability. This enhancement was independent of immediate saccade plans but correlated with the sensitivity to uncertainty in eye movement behavior on longer time scales (across sessions/days). The findings suggest that topographic visual maps receive motivational signals of uncertainty, which enhance the priority of informative stimuli and the likelihood that animals will orient to the stimuli to reduce uncertainty.

## Introduction

Humans and other animals have an intrinsic desire to gain information and predict future events. This desire has been captured in the laboratory using tasks of non-instrumental information demand, in which participants can request advance information about an upcoming reward but cannot use the information to alter the amount of reward^1, 2^. Humans and monkeys sacrifice rewards to obtain non-instrumental information, suggesting that they assign intrinsic utility to information independently of extrinsic incentives^1, 2^.

The studies have also suggested that the intrinsic utility of information is of two kinds. Participants are more likely to demand information when rewards are uncertain (e.g., if a trial has a 50% versus 100% reward probability), suggesting that they are motivated to obtain an early resolution of uncertainty. However, participants are also more likely to demand information when they have high versus low reward probability in the absence of uncertainty (e.g., a 100% versus 0% chance of reward) suggesting that they are motivated to gain positive observations independently of resolving uncertainty. Individual choices reflect weighted combinations of uncertainty reduction and reward drives, suggesting that both motives combine to determine individual information demand^3-5^.

Uncertainty-driven (as opposed to reward-driven) information demand is particularly important because it is theoretically normative and allows animals to maximize predictive accuracy ^4, 6, 7^. However, we have incomplete understanding of its neural mechanisms. In most tasks of information demand, animals resolve uncertainty by making rapid eye movements (saccades) to visual stimuli. Saccades are mediated by fronto-parietal areas that encode the locations of visual stimuli and select saccade goals^8, 9^. Recent reports suggest that these areas are sensitive to uncertainty, but these findings addressed only instrumental conditions, which differ from non-instrumental conditions in that animals obtain information to enhance reward gains^10, 11^. It is unknown how the cells encode saccade plans to resolve uncertainty independently of instrumental incentives^1, 12^.

Encoding of non-instrumental uncertainty about forthcoming rewards is found in subcortical structures including the anterodorsal septum^13^, the basal forebrain^14^ and midbrain dopamine cells^15^. A set of neurons encoding uncertainty during non-instrumental information demand were recently identified in the pallidum, dorsal striatum and the dorsal anterior cingulate cortex (dACC)^16, 17^. However, while these cells encode the motivation to resolve the uncertainty, they are not spatially tuned and do not specify the selection of specific visual stimuli. Thus, a key question is how non-spatial signals encoding the motivation to resolve uncertainty interact with topographic visual maps that specify concrete sampling policies.

Here we examined this question by recording cells in the lateral intraparietal area (LIP), which is implicated in target selection for attention and gaze^9^, in a task of non-instrumental information demand. Monkeys were free (but not incentivized) to reveal an informative visual cue in trials that had a 0%, 50% or 100% prior reward probability, with the 50% condition uniquely inducing reward uncertainty. We show that visuo-spatial activity of LIP cells is modulated by uncertainty independently of reward probability. Uncertainty modulations had distinct latencies and contextual sensitivity relative to those produced by reward probability, and co-varied with the sensitivity to uncertainty in the monkeys’ information seeking saccades. The findings suggest that topographic visual maps receive distinct motivational signals related to non-instrumental information, which enhance visual priority and thus the likelihood that animals will select and process stimuli that reduce the uncertainty.

## Results

### Information seeking is sensitive to reward probability and uncertainty

Two monkeys performed a task of information demand in which they were free to obtain advance information about a trial’s reward (**Fig. 1A**). A trial started with the presentation of a peripheral cue (Cue 1) signaling the prior reward probability (0%, 50% or 100%), followed by a 1 second delay period when the monkeys maintained central fixation (**Fig. 1A**, “Fixation”) and a 2.5 second period in which gaze was unconstrained (**Fig. 1A**, “Free Viewing”). During the first 1.5 seconds of free-viewing, the monkeys had access to a visual mask and could reveal an additional cue (Cue 2) if they held gaze on the mask. The mask then disappeared and, after a 1 second delay, the trial ended with delivery of the outcome – reward or no reward based on the cued probabilities (**Fig. 1A**, “Outcome”).

**Fig. 1.**
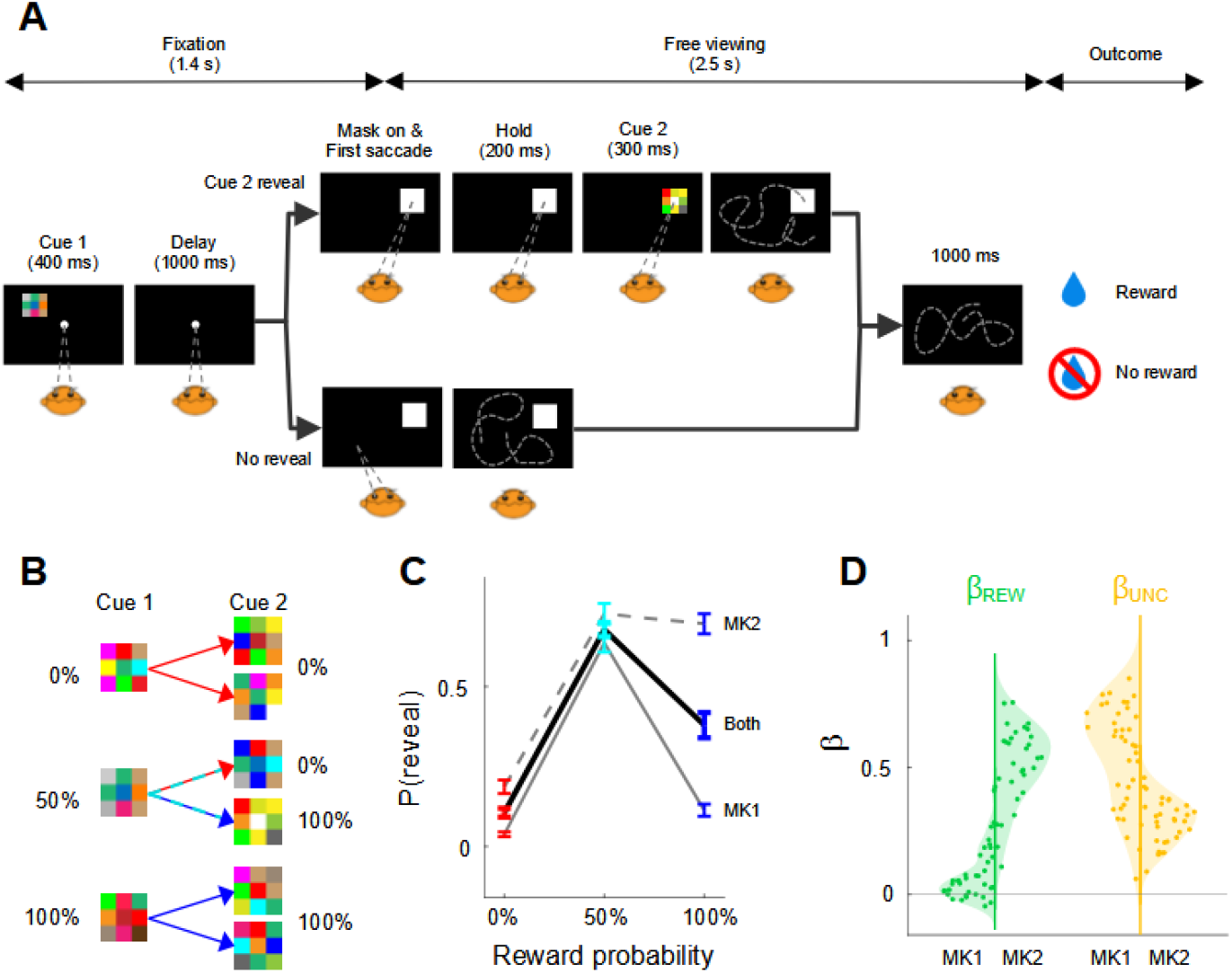
Task and Behavior. **A. Trial structure in the information seeking task** Trials had a period of central fixation (Cue 1 and delay) and a period of free-viewing (mask presentation followed by a blank screen) followed by outcome delivery. During free viewing, monkeys could hold gaze on the mask if they wished to reveal Cue 2. The trial’s outcome (reward or lack of reward) was not contingent on revealing Cue 2. Time intervals (indicated above the individual panels) were fixed, removing uncertainty about the timing of the reward relative to Cue 1 onset. **B. Cue reward contingencies.** Several distinct checkerboards were pretrained to indicate each of three reward probabilities (0%, 50% or 100%). Cue 2 perfectly predicted the trial’s reward. If Cue 1 signaled 0% or 100% reward probability, Cue 2 confirmed this probability. If Cue 1 signaled a 50% probability, Cue 2 was equally likely to signal a reward or lack of reward. Each Cue 1 pattern was equally likely to be followed by one of two Cue 2 patterns at each reward probability. **C. Reveal probability** (P(reveal)) was the fraction of trials where Cue 2 was revealed out of completed trials (in which the monkeys maintained fixation during the initial epoch). Each point shows the mean and SEM. across all neural recording sessions (MK1: n =□37; MK2: n = 31). Here and in all following figures, red, cyan, and blue represent, respectively, 0%, 50%, and 100% reward probability. **D Coefficients indicating the effects of reward probability (β_REW_, green) and reward uncertainty (β_unc_**, **yellow) on information sampling.** Each point is one session. Left and right halves of each distribution show individual monkeys. The shading represents probability density and points are jittered on the x-axis within this envelope for visualization.

A critical feature of the task was that the rewards were non-contingent on free-viewing behavior and the monkeys’ willingness to reveal Cue 2 indicated their intrinsic desire to obtain information. Cue 2, if revealed, provided complete information about the trial’s reward (**Fig. 1B**). Thus, Cue 2 varied in its valence (whether it signaled a reward or lack of reward) as well as the new information it brought relative to Cue 1. On 0% and 100% trials, Cue 2 was redundant, merely confirming the Cue 1 probability, while on 50% reward probability trials, Cue 2 brought new information resolving the prior uncertainty (**Fig. 1B**). Consistent with prior results, the monkeys’ willingness to reveal Cue 2 depended on both factors (**Fig. 1C**). The monkeys were more likely to reveal Cue 2 if they were certain to obtain a reward versus lack of reward (100% vs 0% Cue 1 probability); however, they were also more likely to reveal Cue 2 at 50% relative to 100% reward probability, indicating an additional sensitivity to uncertainty (**Fig. 1C**).

To quantitatively estimate these effects, we fit viewing behavior with a linear model in which one term coded for reward probability (0.0, 0.5 and 1.0) and a second term coded uncertainty (0, 1, 0 for the respective levels of REW; *Methods*, eq. 1). The fitted coefficient for the first term (β_REW_) captured the component of sampling that increased linearly with reward probability, while the coefficient for the second term (β_UNC_) captured the additional role of uncertainty that was not explained by the linear trend. Both β_REW_ and β_UNC_ coefficients were significantly positive in both monkeys on average (β_REW_ relative to 0: MK1: p < 10^-4^, MK2, p < 10^-5^; n = 37 and 31 sessions, respectively; **Fig. 1D**, green; β_UNC_: MK1: p < 10^-6^, MK2, p < 10^-5^; **Fig. 1D**, yellow). β_REW_ coefficients were significant in 37% and 100% of individual sessions in, respectively, MK1 and MK2 and, remarkably, β_UNC_ were significant in 100% of individual sessions in each monkey, indicating a highly consistent sensitivity to uncertainty.

Control analyses ruled out the possibility that these findings had spurious explanations. The monkeys were extensively trained with all the cue patterns (*Methods*) and their anticipatory licking during information demand closely followed the cued reward probabilities (**Fig. S1A**), showing that the monkeys were familiar with and attended to the probabilities signaled by the cues. Second, anticipatory licking was influenced by reward probability even when the monkeys did *not* reveal Cue 2 ruling out that the monkeys mistakenly believed that they had to reveal the cue (**Fig. S1B**, top). Third, viewing behavior was unchanged in control sessions in which the spout was placed inside the monkeys’ mouth, ruling out that the monkeys revealed Cue 2 to reduce the physical effort of licking^3^. Finally, each Cue 1 was equally likely to be followed by one of two Cue 2 patterns (**Fig. 1B**), ruling out that a bias for revealing Cue 2 was related to differential expectations of visual novelty.

### LIP neurons are modulated by reward and uncertainty

To examine the neural substrate of the sensitivity to reward and uncertainty, we recorded the activity of 68 visually responsive neurons (37 in MK 1) from area LIP that is implicated in target selection for attention and gaze. Neurons were tested on the information seeking task if they had well-isolated spike waveforms and spatially tuned delay period activity in a standard memory guided saccade task (**Fig. S2**). We first describe the neurons’ responses to Cue 1 and the mask, followed by analysis of the relationship between neural activity and the monkeys’ saccadic decisions, and the responses to the information conveyed by Cue 2.

During the period of central fixation, the retinotopic locations of Cue 1 and the mask were controlled and could fall inside or outside the receptive field (RF) of the cell (*Methods*). By comparing the effects of reward probability at different stimulus geometries, we could thus examine how the neurons’ visuo-spatial selectivity interacted with their reward or uncertainty selectivity.

As expected from their visuo-spatial selectivity, the neurons had excitatory responses if Cue 1 appeared inside the RF (**Fig. 2A**, left) but not outside the RF (**Fig. 2A**, right) and, upon mask onset, had excitatory responses if the mask appeared inside the RF (**Fig. 2B**, top) but not outside the RF (**Fig. 2B**, bottom). Reward expectations modulated responses at all stimulus geometries. When Cue 1 or the mask were inside the RF, the neurons’ visual responses were higher on trials with 100% and 50% relative to 0% reward probability (**Fig. 2A**, left, and **Fig. 2B**, top), and similar modulations were found when the stimuli were outside the RF (**Fig. 2A**, right and **Fig. 2B**, bottom).

**Fig. 2.**
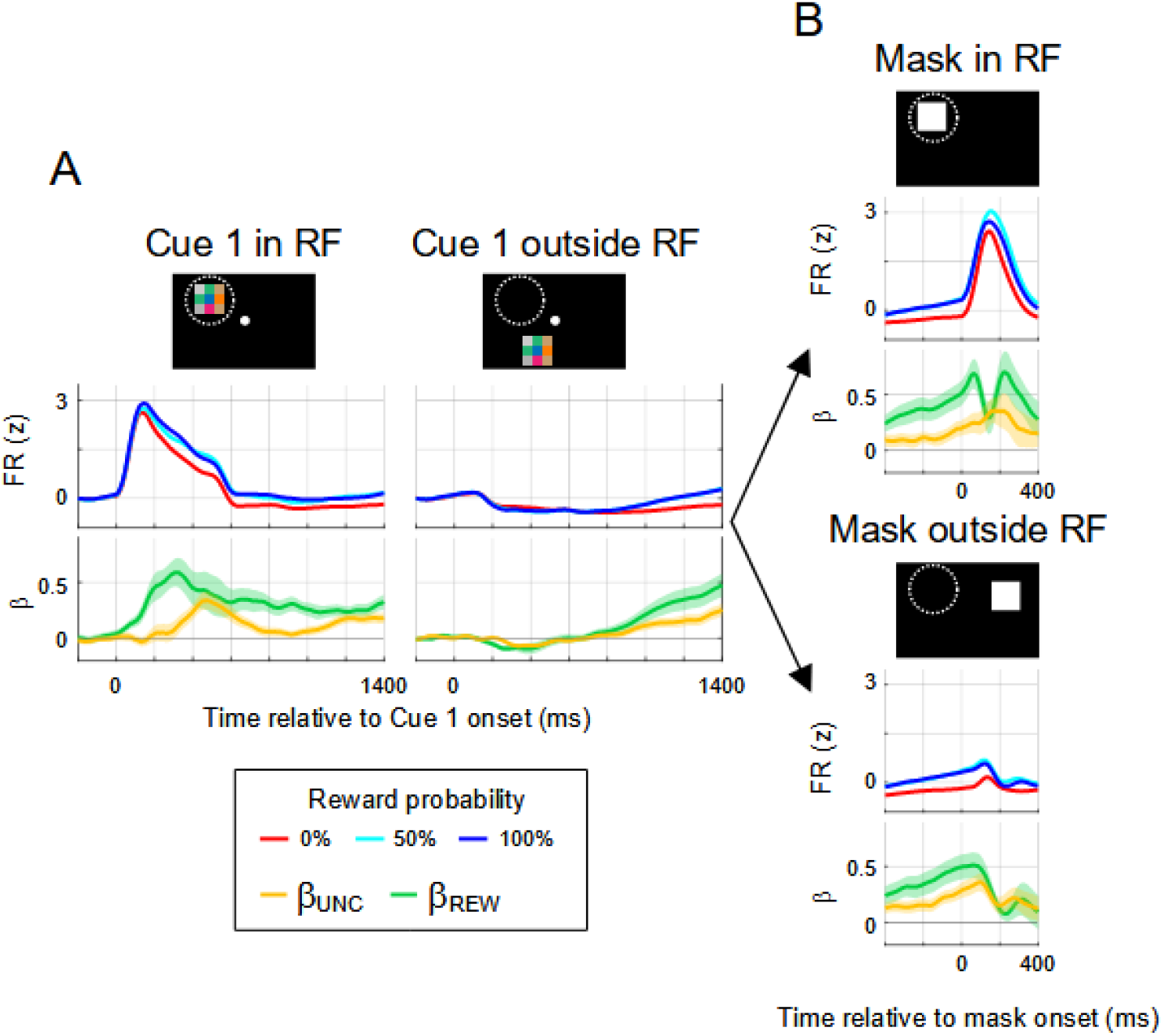
LIP modulations by reward and uncertainty. **A. Responses aligned to Cue 1 onset.** The cartoons show Cue 1 (colored checkerboard) appearing inside or outside the RF (dashed circle). The middle row shows population peri-stimulus response histograms computed by z-scoring the raw FR within each neuron (using all of its data in the task), averaging across trials within each neuron, and averaging across neurons (n = 68). The bottom panels show regression coefficients indicating the effects of uncertainty (β_UNC_, yellow) and reward (β_REW_, green). The traces are averaged across cells (n = 68) and shading shows 2 SEM (equivalent to 95% confidence interval) **B. Responses aligned to mask onset.** Same format as in A. Saccades toward or away from the mask typically occurred in the first 200 ms (average latency 193.2±0.22 ms).

To measure the contributions of uncertainty and reward probability to these modulations, we fit firing rates with a model that included the reward and uncertainty regressors we used for behavior alongside regressors indicating the direction and latency of the first saccade away from fixation, the prior trial reward, and licking behavior (*Methods*, eq. 3). The fitted β_REW_ and β_UNC_ coefficients thus indicated the effects of reward and uncertainty independently of other factors that may have affected neural activity. The β_REW_ coefficients captured a linear coding of reward probability while the β_UNC_ coefficients indicated an effect of uncertainty beyond that predicted by the linear trend.

After onset of Cue 1 inside the RF, β_UNC_ and β_REW_ coefficients were positive indicating that activity was independently enhanced by reward probability and uncertainty (**Fig. 2A**; left). Both effects were sustained and remained significant throughout the end of the delay period (1000-1400 ms; β_UNC_: 0.14±0.02, p < 10^-9^, β_REW_ 0.25 ± 0.02, p < 10^-11^, Wilcoxon test relative to 0, n=68). Similarly, after onset of the mask inside the RF, both coefficients were significantly greater than 0 (**Fig. 2B,** top; 0-400 ms; β_UNC_: 0.25 ± 0.04, p < 10^-6^; β_REW_: 0.51 ± 0.05, p < 10^-10^, n=68).

When Cue 1 was presented outside the RF, the neurons showed an initial slight suppression followed by a low-level response anticipating the possible onset of the mask inside the RF (**Fig. 2A**, right). This latter response was significantly modulated by reward and uncertainty (1000-1400 ms, β_UNC_: 0.16±0.02, p < 10^-9^; β_REW_: 0.34±0.04, p < 10^-9^, n=68). Similarly, when the mask appeared outside the RF, firing rates were significantly modulated by reward and uncertainty (**Fig. 2B**, bottom; βUNC: 0.23±0.03, p < 10^-8^; β_REW_: 0.27±0.03, p < 10^-8^, n=68). βUNC and β_REW_ coefficients after inside- and outside-the-RF presentations had comparable magnitudes (paired tests, Cue 1 β_UNC_, p = 0.33, Cue 1 β_REW_, p = 0.12; mask β_UNC_: p = 0.23) although β_REW_ coefficients were higher after mask onset inside versus outside the RF (β_REW:_ p < 10^-6^, n = 68). The significant modulations at non-visual (blank) locations seem distinct from the more localized reward modulations found in operant tasks^18, 19^, and this difference may be due to the absence of operant training and/or competing stimuli in our task.

Because the reward and uncertainty modulations co-existed throughout the task epochs, a possible concern is that the neurons simply encoded the *possibility* of reward – responding equivalently to 50% and 100% trials rather than separately to reward and uncertainty. Three analyses argue against this interpretation. First, a formal model comparison showed that replacing the separate reward and uncertainty terms with a single term of “reward possibility” (i.e., where trials were coded as 0 if they had 0% probability and 1 otherwise) reduced model fit for most cells (average increase in AIC score of 3.80±1.01).

Second, uncertainty modulations had significantly longer latencies relative to reward modulations even when controlling for relative signal strength. When Cue 1 was inside the RF (**Fig. 2A**), the average latency for significant selectivity for β_UNC_ coefficients was 283 ± 31 ms versus 174 ± 22 ms for β_REW_ coefficients (Wilcoxon paired test, p < 0.01; n = 67 cells with a detectable latency for each signal). To rule out that apparently longer latencies resulted from weaker signal strength (and potentially lower statistical power for detecting uncertainty modulations), we repeated the analysis in random subsamples of cells that were matched in their peak β_UNC_ and β_REW_ coefficients (*Methods*). Across 1,000 subsamples, the latency of uncertainty modulations was longer by a median of 53 ms with a 95% confidence interval that did not include 0 ([96, 22] ms), confirming that uncertainty selectivity arose significantly later than reward selectivity at equivalent signal strength.

A third line of evidence establishing the independence of uncertainty and reward modulations came from a subset of cells that were also tested in a passive task in which the Cue 1 stimuli were presented while the monkeys maintained passive fixation (**Fig. S3**). If the cells simply encoded the possibility of reward, responses to 50% and 100% probability should be similar both contexts. However, while responses to 100% stimuli were similar in the passive and active sampling conditions, responses to 50% stimuli were much lower in the passive condition (**Fig. S3A**). This produced, in the passive condition, a negative modulation by uncertainty (**Fig. S3B**) that was starkly distinct from the positive effects of reward probability (**Fig. S3C**). Thus, uncertainty modulations in LIP cells differed from those produced by reward probability in their dynamics and contextual modulations.

### Neurons independently encode uncertainty and saccade plans

The fact that β_UNC_ and β_REW_ coefficients were estimated after controlling for saccade direction and latency suggests that these factors modulated visual responses independently of saccade motor plans. Indeed, the analyses provided little evidence that the cells encoded saccade motor plans after Cue 1 presentation or when the mask appeared outside the RF. For these geometries, the coefficients for saccade direction and latency were not statistically significant and follow up analyses showed no consistent effects of the decision to reveal Cue 2 independently of directional selectivity.

However, consistent with the fact that saccade motor plans modulate the neurons’ visual activity^20^, the cells did have a saccadic response if the mask was inside the RF, when they had higher firing rates if the saccade was directed toward versus away from the RF (**Fig. 3A**, top). We thus examined how these saccade modulations were combined with the effects of uncertainty and reward. One possibility is that the cells would show interactive effects, encoding saccade planning only under specific levels of uncertainty or reward probability as has been reported for basal ganglia/dACC cells^16^. Alternatively, the cells may encode saccade plans and reward/uncertainty as independent responses.

**Fig. 3.**
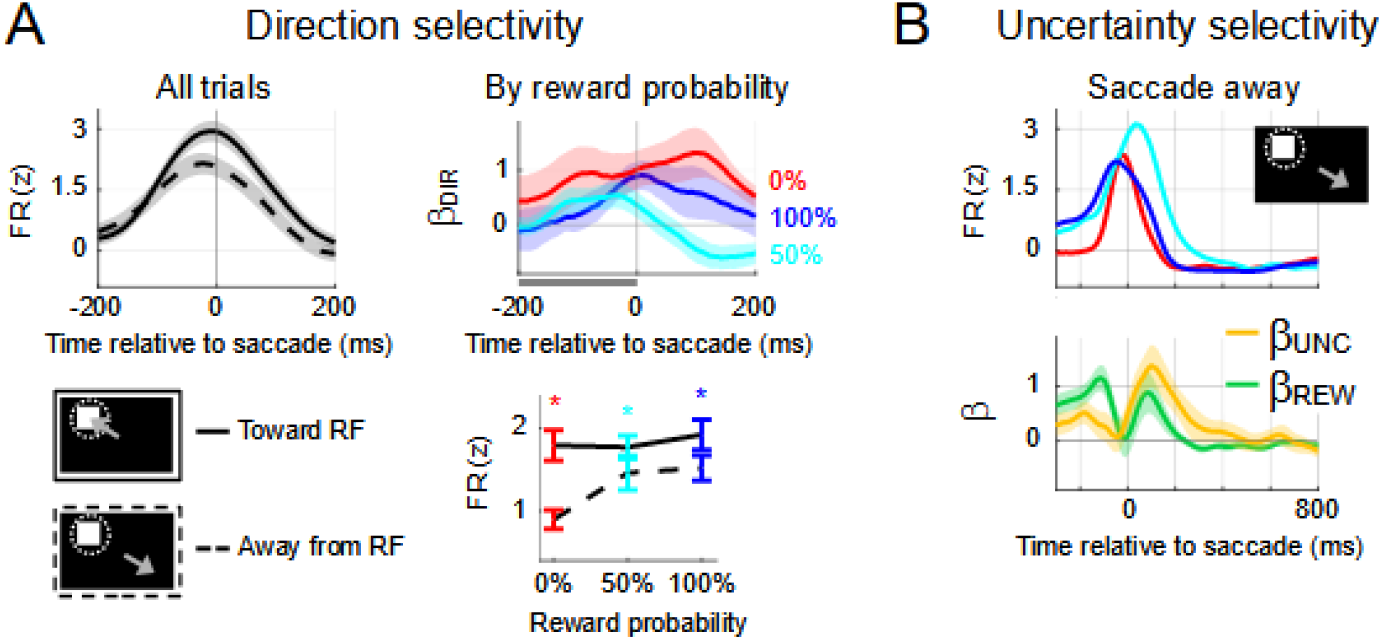
Uncertainty and saccade direction. **A. Directional selectivity.** (Left) PSTH of activity aligned on saccade onset, pooled across reward probabilities for the geometry depicted in the cartoons (mask inside the RF and saccades toward (solid) and away (dashed) from the RF). The traces show mean and 2 SEM for all cells with sufficient trials in each condition (saccade away, n = 68; saccade toward, n = 52). (Right) **Directional selectivity by reward probability.** Top: Time-resolved regression coefficients capturing the effect of saccade direction (β_DIR_) in the trials in A separated by reward probability (mean and 2 SEM across cells with sufficient trials; 0%; n = 31; 50%: n = 26; 100%: n = 21). Bottom: Firing rates in the 200 ms before saccade onset for each reward probability and saccades toward (solid) and away from the RF (dashed). Asterisk indicates p<0.05 in post-hoc signed rank tests. **B. Effects of reward and uncertainty for saccades away from a mask in the RF (cartoon).** PSTHs (top) and reward and uncertainty coefficients (bottom) aligned on saccade onset (average and 2 SEM, n = 26 cells with sufficient number of trials). Other conventions as **in Figure 2**.

The findings supported the latter hypothesis. In one analysis, we computed directional selectivity separately for each level of reward probability (**Fig. 3A**, right). Directional coefficients were significant in all cases and, in the 100 ms before the saccade, had no significant effect of reward probability (p = 0.087, 1-way ANOVA; n = 31, 26 and 21 cells for, respectively, 0%, 50% and 100% probability). A follow up analysis on raw firing rates confirmed this conclusion (**Fig. 3A**, bottom right), with a 2-way ANOVA showing a significant effect of saccade direction (p < 10^-3^), but no effect of reward probability or interaction between saccade direction and probability. With the exception of low firing rates for saccades away at 0% probability, reward probability did not differentially affect firing rates before saccades toward the RF (stars).

A second analysis confirmed this conclusion by showing that uncertainty modulations were not predicated on a specific saccade. If the mask was inside the RF but the saccade was away and did not reveal Cue 2, responses at 50% were nevertheless significantly higher than those at 100% reward probability, indicating a highly robust uncertainty modulation (**Fig. 3B**, top). β_UNC_ and β_REW_ coefficients were statistically significant before the saccade (-200 to 0 ms; β_UNC_, 0.38±0.09, p < 0.01; β_REW_ = 0.57±0.09, p < 10^-4^; n = 26 cells) and after saccade onset (0-200 ms; βUNC, 1.06±0.15, p < 10^-4^; β_REW_ = 0.54±0.14, p < 10^-3^). Thus, the cells encoded uncertainty independently of the trial by trial saccade or reveal decision.

### Neural and behavioral uncertainty modulations correlate across days

Given the independence of the uncertainty modulations from the immediate saccade plan, we asked if these modulations correlate with the monkeys’ sensitivity to uncertainty on longer time scales – i.e., across sessions rather than individual trials. To examine this question, we again focused on trials in which the monkeys did *not* reveal Cue 2, ensuring that we capture correlations that are independent of the immediate information seeking decision.

The effect of uncertainty on the monkeys’ revealing behavior showed daily variability (cf **Fig. 1D**) and that was correlated with the uncertainty modulation of the LIP visual response to the mask. This was readily appreciated by plotting the neural β_UNC_ coefficients for all cells in order of the behavioral β_UNC_ coefficients (**Fig. 4A**). When the mask was inside the RF and the monkeys showed high β_UNC_ coefficients in their revealing behavior (**Fig. 4A**, left, top rows), the visual response to the mask showed pronounced enhancement by uncertainty, which lasted for several hundreds of milliseconds even though the saccade was away from the mask (cf. **Fig. 3B**). This resulted in a highly significant correlation between neural and behavioral β_UNC_ coefficients that was consistent in both monkeys (Spearman correlation, combined data set, rho = 0.46, p < 0.001, n = 59; MK1, rho = 0.41, MK2, rho = 0.55, both p < 0.05). In contrast, when the mask was outside the RF, correlations were weaker and significant in only one monkey (**Fig. 4A**, right; rho = 0.35, p < 0.01, n = 66; MK1, rho = 0.39, p < 0.05, MK2, rho = 0.30, p = 0.12), showing that the effect was strongest for the mask-evoked visual activity.

**Fig 4.**
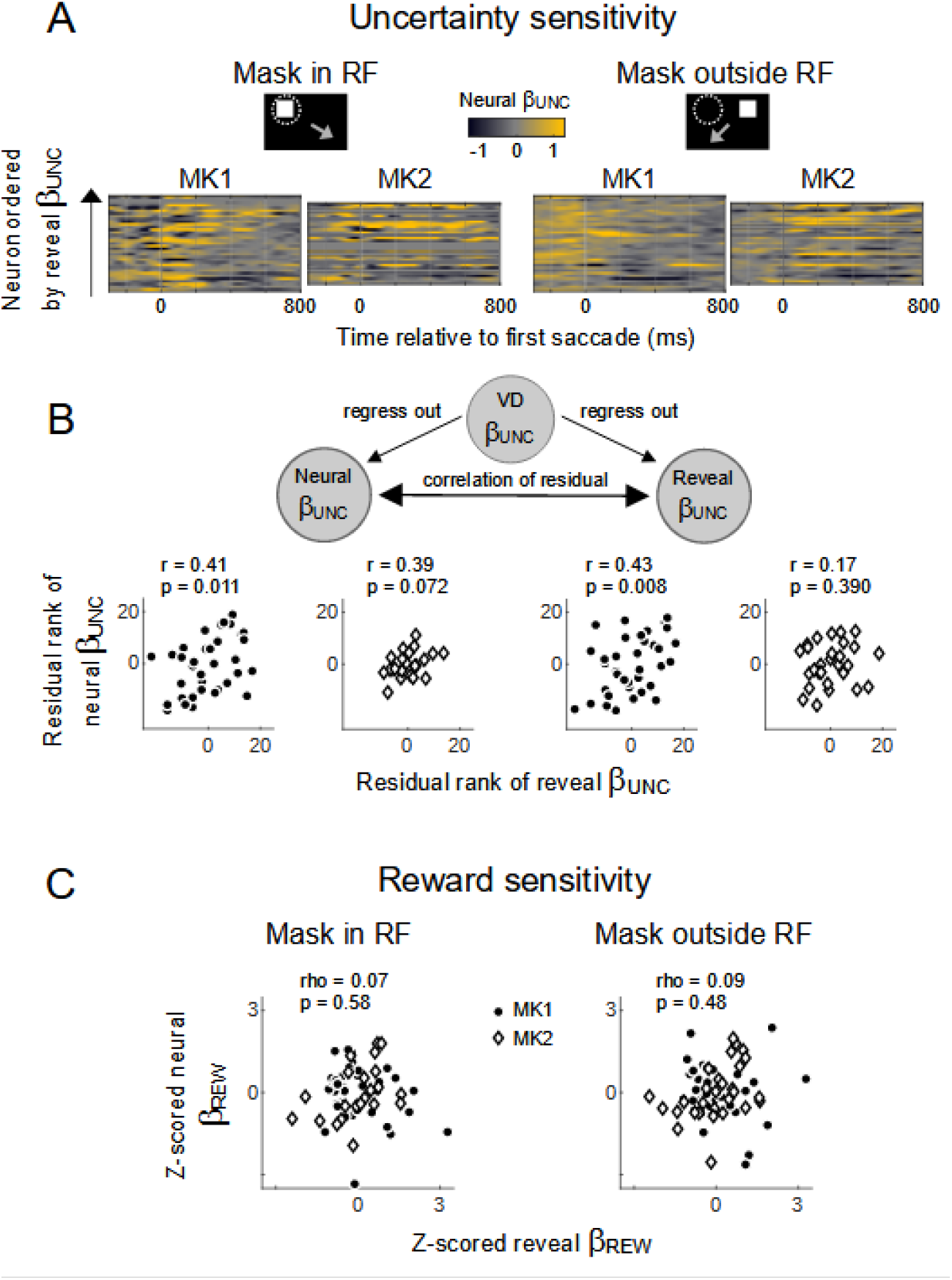
Correlations between neural and behavioral sensitivity to uncertainty. **A.** Color maps of the time-resolved β_UNC_ coefficients (color) for individual neurons ordered according to the magnitude of the uncertainty coefficient in revealing behavior (reveal β_UNC_). The analysis was performed on trials when the mask was inside (left) or outside the RF (right) but the saccade was away from the mask (cartoons). **B.** Top: the partial correlation analysis controlling for correlations between the effects of uncertainty on VD and reveal behavior (see text for details). Bottom: The scatter plots show the correlations of residual rank corresponding to the panels in A. Each point shows the residual rank of the neuronal β_UNC_ coefficient 200-600 ms after saccade onset (ordinate) against the residual rank of the reveal β_UNC_ coefficient in the same session (abscissa). The text shows the correlation coefficient and its p-value. **C.** Correlations for neural and reveal β_REW_ coefficients (combined across monkeys after z-scoring within monkey to remove individual effects).

The post-saccadic time course of these correlations raises the possibility that they may encode some aspect of the post-saccadic processing of Cue 2 rather than revealing behavior. Indeed, when the monkeys revealed Cue 2, their viewing durations (VD) were longer if Cue 2 signaled a positive versus negative outcome (an effect of reward) and if Cue 2 resolved uncertainty rather than confirming prior expectations (an effect of uncertainty). Analysis of VD with a model including reward and uncertainty regressors (*Methods*, eq. 2) showed that the uncertainty coefficients (β_UNC_VD_) were significant and, across days, were positively correlated with the β_UNC_ coefficients on reveal probability (MK1, rho = 0.43, p < 0.01; MK2, rho = 0.39, p < 0.05). Thus, the neurons may encode a variable impact of uncertainty on VD that showed persist even on no-reveal trials.

We thus carried out a non-parametric partial correlation analysis to rule out spurious effects of VD uncertainty modulations (**Fig. 4B**, cartoon). We calculated the ranked coefficients for each variable (rβ), regressed out the effects of VD on neural activity (by fitting rβ_UNC_neural_ ~ 1 + rβ_UNC_VD_) and the effects of VD on reveal probability (by fitting rβ_UNC_reveal_ ~ 1 + rβ_UNC_VD_) and examined the residuals of the β_UNC_neural_ and β_UNC_reveal_ coefficients. The correlations of the rank residuals remained significant in each monkey (**Fig. 4B,** scatterplots). This effect was replicated when we included reveal trials (**Fig. S4**) although this analysis was less reliable because on these trials the neurons also responded to the information by Cue 2 (described in the following section). Thus, the effect of uncertainty in LIP neurons was correlated with the effects of uncertainty on reveal probability independently of a relationship with VD.

In contrast to these findings in the response to the mask, we found no equivalent correlations in the visual/delay responses that were evoked by Cue 1 or those anticipating mask onset. Importantly, we found no correlations between the neural and behavioral *reward* sensitivity in any task epoch, including when the mask was inside or outside the RF (**Fig. 4C**, inside RF: rho = 0.07, p = 0.58; opposite RF: rho = 0.09, p = 0.48; n = 68 cells, combined after z-scoring within monkeys). In sum, daily variability in the effect of uncertainty on revealing behavior correlated with the effect of uncertainty on the LIP responses to informative stimuli independently of saccade motor plans. The correlations were specific to uncertainty rather than reward modulations and were manifest even when cells were independently sampled across days, suggesting that they are global and uniform across LIP cells.

### The neurons respond to new information conveyed visually

After the initial saccade away from fixation, the monkeys had a period of free-viewing and, in some trials, revealed Cue 2. We found that, when the monkeys revealed Cue 2, the neurons encoded the reward and uncertainty signaled by the cue even though the RF was no longer aligned with the cue. However, the cells did not respond on no-reveal trials, when the monkeys anticipated the resolution of uncertainty by the outcome itself.

When the monkeys revealed Cue 2, many neurons showed a transient decline in firing if Cue 2 announced a lack of reward but no change if Cue 2 announced a reward, showing that they were transiently enhanced by the reward announced by Cue 2 (**Fig. 5A**, top panel, red vs blue). Moreover, the firing rate drop on no reward trials was more pronounced if Cue 2 resolved uncertainty rather than merely confirming prior expectations, indicating a consistent integration of reward and uncertainty (**Fig. 5A**, top, cyan-red vs solid red traces).

**Fig. 5.**
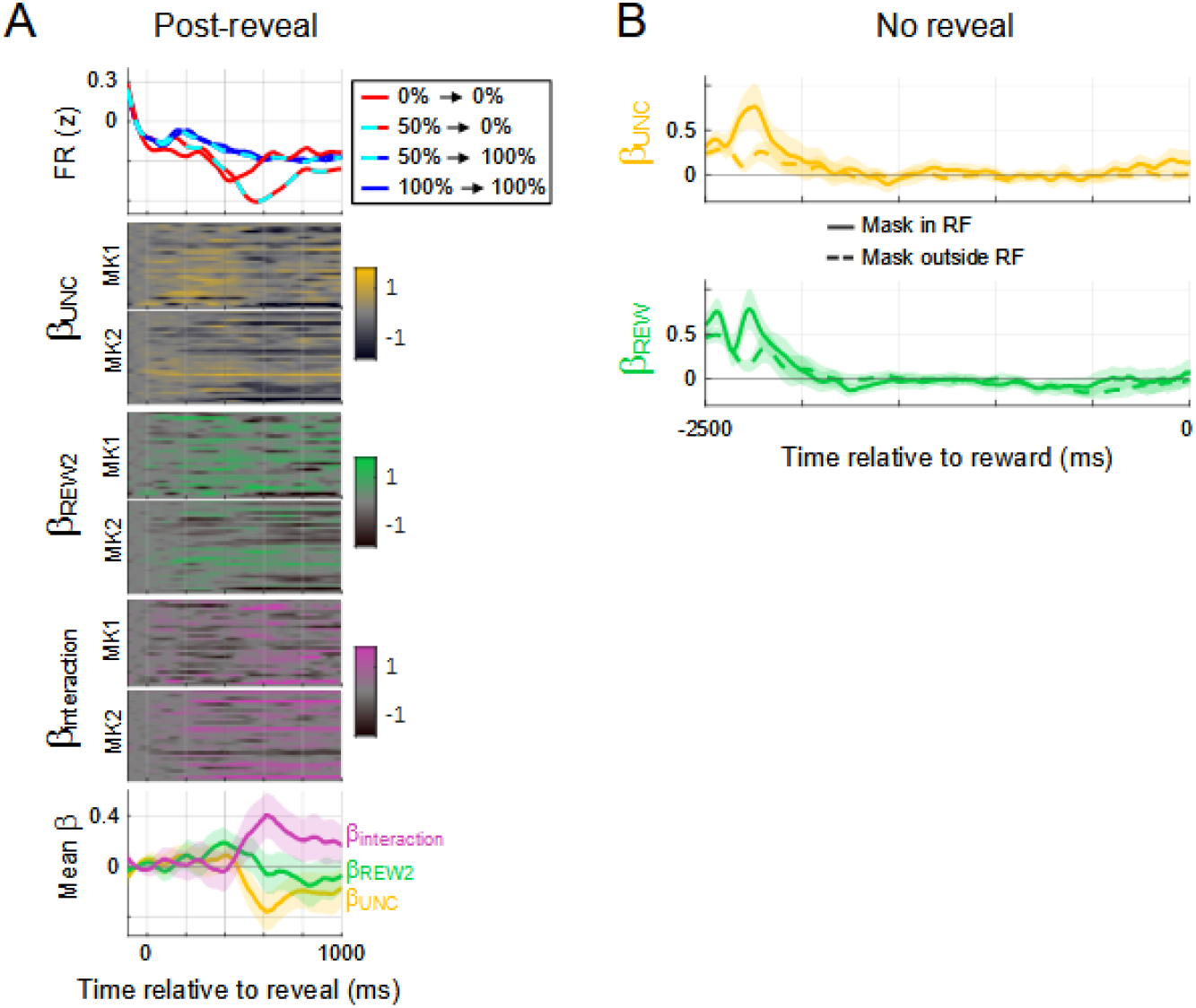
LIP neurons responded to new information. **A.** Top: Firing rates on trials in which Cue 2 was revealed, aligned on the time of reveal. PSTHs show average firing for the possible combinations of reward and uncertainty resolved by Cue 2 (n = 60 cells with sufficient number of trials). Pre-reveal firing was subtracted to remove effects of Cue 1. Heatmaps show time-resolved regression coefficients (uncertainty, reward and interaction) for each neuron included in the top panel. The bottom panel shows the average coefficients (shading shows 2 SEM).

We quantified these effects using a linear model with regressors indicating the reward signaled by Cue 2 (REW2, 0 or 1), the uncertainty resolved by Cue 2 (UNC, as for Cue 1) and the interaction of REW2 x UNC (we subtracted pre-reveal firing rates to remove the effects produced merely by Cue 1; *Methods*, eq. 4). The uncertainty coefficients peaked at ~600 ms after revealing Cue 2, were *negative* (showing that uncertainty suppressed firing for Cue 2 signaling no-reward) and were seen in many individual cells (**Fig. 5A**, top colormap; note the darker hues around 600 ms). The reward coefficients were positive (as firing rates were higher for 100% relative to 0% Cue 2 probability) and were mostly captured by a *positive* interaction with reward (showing that this suppression enhanced the positive effect of reward; **Fig. 5A**, pink).

These effects could not be attributed to coincidental effects of free viewing. The effects were independent of the direction and latency of the saccade to the mask, the prior trial reward, lick rate and viewing duration, which were included as nuisance regressors (*Methods*, eq. 4). The effects peaked after Cue 2 was covered again by the mask (300 ms after the reveal) and, across days, did not correlate with the uncertainty or interaction coefficients on VD. Similarly, the effects were not mere passive continuations of a mask-related response since the effects of reward or uncertainty before and after the reveal were not correlated. Finally, the dip in activity on no-reward trials did not reflect a coincidental decrease in the fraction of preferred free-viewing saccades; the fraction of saccades with vectors similar to (within 45 degrees of) the cells’ preferred vector was *higher* when Cue 2 announced a lack reward versus a signaling a reward (41.9%±1% vs 32.6%±1% of all saccades; n = 68 sessions).

On *no-reveal trials*, the monkeys could anticipate that reward uncertainty would be resolved by the outcome itself. However, the cells did not encode uncertainty or reward in this epoch, whether the mask had been inside or outside the RF (**Fig. 5B**), suggesting that their sensitivity to these factors was pronounced when reward information was signaled visually but not by the actual outcome.

## Discussion

The vast majority of empirical studies of selective attention have relied on instrumental tasks in which participants discriminate sensory stimuli to guide an incentivized choice (e.g., a monkey is trained to attend to a Gabor patch to correctly report its orientation and receive a reward^21^). Consistent with this empirical focus, instrumental rewards have been shown to modulate fronto-parietal areas and proposed to contribute to guiding attention to choice-relevant stimuli(e.g., ^22, 23^).

However, decision-theoretic frameworks, supported by recent studies of non-instrumental information demand, propose that attention is more fundamentally motivated by the *uncertainty* that can be resolved by a stimulus independently of instrumental incentives^1, 6^. Our findings reveal a cellular mechanism by which uncertainty modulates visual salience independently of instrumental and non-instrumental incentives. We show that uncertainty modulated visual responses in LIP cells, and these modulations had distinct latencies and contextual selectivity relative to those produced by reward probability. While uncertainty modulations did not predict trial by trial saccades, they enhanced the visual responses to informative stimuli in a manner that was correlated, over stretches of trials, with the probability that monkeys would make saccades to those stimuli. The findings suggest that LIP cells combine visual responses with motivational signals related to resolving uncertainty and prioritize the selection of stimuli that reduce the uncertainty.

Several aspects of our findings suggest that uncertainty information is conveyed to LIP from neurons without visuo-spatial selectivity. Upon onset of Cue 1 and the mask, uncertainty modulated responses to visual stimuli and at non-stimulated locations. After a saccade away from fixation, uncertainty had long lasting effects, whether the saccade was away from the mask or was toward the mask and revealed Cue 2. Finally, neural activity was significantly correlated with behavior although neurons were independently sampled across days, suggesting that uncertainty had relatively uniform effects on populations of LIP cells.

Untuned signals of uncertainty may reach LIP through multiple pathways. Some of these pathways may involve neuromodulators, including midbrain DA neurons that encode reward uncertainty^15^ and can affect LIP directly or through the frontal eye field^24^, or acetylcholine and norepinephrine, which have been implicated in uncertainty, executive function, and modulations of visual activity^25, 26^. A particularly interesting hypothesis is that LIP interacts with a newly-described network of information demand comprising neurons in the pallidum, dorsal striatum and dACC^17^. Similar to our results, a subset of cells in these structures respond more for uncertain relative to certain (100%) prior reward probabilities and predict the monkeys’ willingness to make saccades that resolve non-instrumental uncertainty. These neurons may be part of a distributed network that generates the motivation to resolve uncertainty (through the basal ganglia/dACC circuit) and specifies concrete active sensing/attentional policies for obtaining this goal (through the LIP visual map).

Comparing our results with those in the basal ganglia/dACC provides clues about the computations carried out by this network. Unlike the visuo-spatial tuning of LIP cells, basal ganglia/dACC cells were not spatially tuned and anticipated the timing but not the locus of the resolution of uncertainty^17^. Moreover, while LIP neurons reported only the resolution of uncertainty through *visual* stimuli, basal ganglia/dACC cells anticipated uncertainty resolution through both visual stimuli and reward delivery (tactile/gustatory/auditory modality). Thus, the basal ganglia/dACC circuit seems to provide supra-modal signals indicating the time of uncertainty resolution, while LIP encodes attentional signals specific to the sensory modality that resolves the uncertainty.

The two circuits also differed in their integration of information about gaze and uncertainty. Neurons in the basal ganglia/dACC circuit predicted the time of gaze shifts only in the presence of uncertainty – at 50% but not 0% or 100% reward probability^17^ – whereas LIP cells showed equivalent pre-saccadic enhancement at all levels of uncertainty and reward probability (**Fig. 3**). Thus, the basal ganglia/dACC may interact with oculomotor structures according to motivational state, with different subsets of cells being recruited if saccades aim to resolve uncertainty versus viewing conditioned stimuli, whereas LIP integrates motivational factors into a common signal of priority for attention and gaze^9^.

Our finding that uncertainty modulated LIP *visual* rather than saccadic responses is consistent with the view that this area prioritizes visual stimuli, while saccadic motor decisions are made in downstream structures (e.g., the frontal eye field or superior colliculus)^19, 27^. A recent result from our lab suggests that one role of an uncertainty-modulated visual representation may be to coordinate the *selection* of an informative stimulus with the post-saccadic *processing* of the stimulus. We showed that, in a task of instrumental information demand, uncertainty-related enhancements of LIP activity before the saccade correlated with the monkeys’ efficiency in making decisions based on the information after the saccade^10^. Similarly, in the present non-instrumental conditions, we find that the cells encoded both the prioritization of an informative stimulus and the reward and uncertainty that were signaled by the stimulus. Interestingly, in both the present and earlier investigations^10, 28^, we found little evidence that LIP neurons encode the monkeys’ dwell time on the revealed information, consistent with abundant evidence the timing of saccades (toward or away from stimuli) is controlled by separate circuits^29, 30^. However, our results suggest that the cells tag visual stimuli across a saccade and, while selecting a visual stimulus, may also prepare the brain for extracting and using the information that will be gleaned from the stimulus – an important hypothesis for future investigations.

## Methods

### General

Data were collected from two adult male rhesus monkeys using standard behavioral and neurophysiological techniques^31^. All methods were approved by the Animal Care and Use Committees of Columbia University and the New York State Psychiatric Institute as complying with the guidelines within the Public Health Service Guide for the Care and Use of Laboratory Animals. Behavioral control was implemented in MonkeyLogic, stimuli were presented on a Mitsubishi Diamond Pro 2070 monitor (30.4×40.6 cm viewing area), eye tracking was performed using an Applied Science Laboratories model 5000 (digitized at 240Hz), licking was recorded with an in-house device that detected interruptions in a laser beam produced by extensions of the monkeys’ tongue, and action potentials were recorded using an APM digital processing module (Fred Haer). Individual electrodes (glass-coated tungsten electrodes,

Alpha Omega, impedance at 1 kHz: 0.5–1MΩ) were inserted in daily sessions and aimed to the lateral bank of the intraparietal sulcus based on stereotactic coordinates and structural magnetic resonance imaging.

### Memory Guided Saccade Task

After obtaining a well-isolated waveform, a neuron was first screened with a standard MGS task in which a peripheral target was flashed for 300 ms while the monkeys maintained central fixation and, after a 500 ms delay period, the monkeys were rewarded for making a saccade to the remembered target location. Neurons were further tested only if they had spatially tuned visual and delay period responses on this task (**Supplementary Fig. 2**). For these cells, the RF was mapped by conducting the MGS at the same locations used in the information-seeking task, including the estimated RF center and two equally eccentric locations spaced at 120° intervals.

#### Information sampling task

The monkey fixated on a central point to initiate a trial. A pattern (Cue 1) indicating 0%, 50%, or 100% reward probability then appeared for 400 ms, followed by a 1,000 ms delay period and the onset of a white mask concealing Cue 2. The locations of Cue 1 and Cue 2 were randomly selected from 3 possible equi-eccentric and equidistant locations, with the constraint that they did not overlap in a trial. The fixation point was removed simultaneously with mask onset and the monkeys were free to deploy gaze for 2,500 ms. For the first 1,500 ms of this epoch, the mask remained visible and the monkeys could reveal Cue 2 by fixating the mask for a minimum of 200 ms (ensuring that their gaze did not spuriously land on the mask). If revealed, Cue 2 was visible for 300 ms and was again concealed by the mask regardless of the monkey’s gaze location. The mask then disappeared and, after a 1,000 ms blank screen, the trial ended with a tone and the delivery of the outcome - reward or no reward - as predicted by the cues regardless of free-viewing behavior. All temporal intervals between Cue 1 onset and outcome were fixed, removing uncertainty about the delay to reward delivery.

Before experiencing the information seeking task, the monkeys were extensively familiarized with 8 cue patterns - square colored checkerboard (“Mondrian”) measuring 3 deg of visual angle that were equated for luminance and discriminability^32^. Three patterns were consistently associated with 0% reward probability, two with 50% probability and three with 100% reward probability. Each trial was first assigned a reward probability and outcome (reward/no-reward) each randomized with uniform probability across trials. Then, one of the patterns signaling the appropriate probability was randomly assigned as Cue 1 with each Cue 1 pattern followed by two equiprobable Cue 2 patterns signaling the appropriate outcome.

### Passive viewing task

On each trial, the monkey maintained fixation on a central point, viewed a Cue 1 pattern that was presented for 300 ms in the periphery and, after an additional 1,100 ms fixation period, received a reward with the probability assigned to the cue. Trials were run in blocks of two types, one containing only 50% Cue 1 and the other containing 0% and 100% Cue 1 which were randomly interleaved.

### Data analysis

During neural recordings, one of the 3 possible locations was in the RF of the cell while the others were outside the RF. We analyzed data from completed trials in which the monkeys successfully maintained fixation and either did not reveal Cue 2 or did so within 600 ms of mask onset (on average, 607 trials per cell; for simplicity, we excluded < 0.5% of trials in which Cue 2 was revealed after more than 600 ms).

#### Behavior

Saccade onset and offsets were detected based on velocity and acceleration criteria using custom-made software^19^. To analyze information seeking behavior, we fit each session’s data to a linear model where REV = 1 if Cue 2 was revealed and 0 otherwise, REW is the reward probability signaled by Cue 1 (0, 0.5, or 1), and UNC is the associated uncertainty (0, 1, 0).

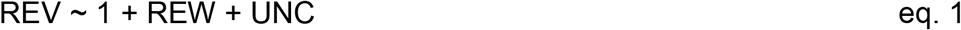

To analyze viewing duration (VD), we fit the reveal trials in each session to a linear model:

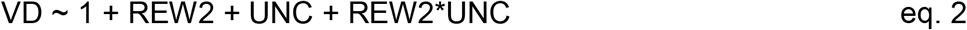

where VD is the time from removal of the mask and eye’s exit from a 2 deg window surrounding the mask, REW2 is the reward probability signaled by Cue 2 (0 or 1) and UNC is the uncertainty resolved by Cue 2 defined as above.

#### Neural analysis

We computed z-scored firing rates (FRz) by convolving each trial’s spike trains with a Gaussian filter (sigma = 30 ms) and z-scoring within a cell using all the time points and trials collected for that cell. We analyzed each cell using statistical models as noted below, and report coefficient distributions over all the cells that had at least 2 trials for each condition required to estimate the model regressor.

To extract the time-resolved effects of reward and uncertainty in the information seeking task (**Fig. 2-5**), we fit FRz with 1 ms resolution throughout the period of interest using the equation:

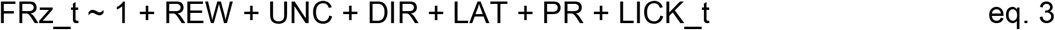

where FRz_t is the z-scored firing rate at time t, REW and UNC are defined as in eq. 1, DIR is the direction of the first free-viewing saccade (1 if directed in a ±45 degree cone centered on the RF; 0 otherwise), LAT is the latency of the first free-viewing saccade, PR is the reward outcome on the preceding trial (1 if rewarded, 0 otherwise), LICK_t is the binary licking status at time t (1 if licking, 0 otherwise).

For the passive task, we used the same equation while omitting the DIR and LAT regressors.

To estimate response latencies, we obtained a null distribution (by randomly shuffling REW and UNC trial labels and repeating the regression procedure for 1,000 iterations). We defined signal onset as the first of 3 consecutive milliseconds in which the true coefficient was more extreme than 95% of the null distribution (p < 0.05) and verified that the results held for a range of criteria. To control for the fact that β_REW_ were slightly larger than β_UNC_, we calculated the latency difference between the reward and uncertainty modulations in 1,000 random subsamples of 42 cells in which the distributions of peak β_REW_ and peak β_UNC_ were identical (identical numbers of cells in each magnitude bin, confirmed at several bin sizes).

In the post-reveal analysis (Fig. 5), in order to control for effects of reward and uncertainty merely based on Cue 1, we first subtracted the mean over the 100 ms before Cue 2 onset from the z-scored firing rate on each trial. We then used these mean-subtracted firing rates (FRzd_t) to estimate the effects produced specifically by Cue 2 by fitting the equation:

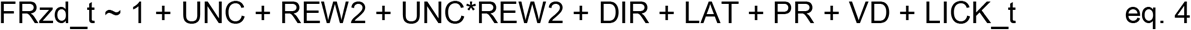

where UNC, DIR, LAT, PR and LICK_t are defined as in eq. 3 (with DIR and LAT referring to the first saccade away from fixation, which triggered the reveal) and VD defined as in eq. 2.

## Acknowledgments

The research described in this paper was supported by a National Eye Institute Grant R01EY025158 to JG, a National Institute of Mental Health R01 MH-098039 to JG, scholarships from Knud Højgaards Fond, Reinholdt W. Jorck og Hustrus Fond, and Viet-Jacobsens Fond to MH. We thank Xue-Xin Wei and Nicholas Foley for their generous help with early stages of data analysis.

## Contributions

ND and JG designed the experiment, ND collected the data, ND, YL and MH analyzed the data, JG and YL wrote the manuscript.

---

**Supplementary Figure 1.**
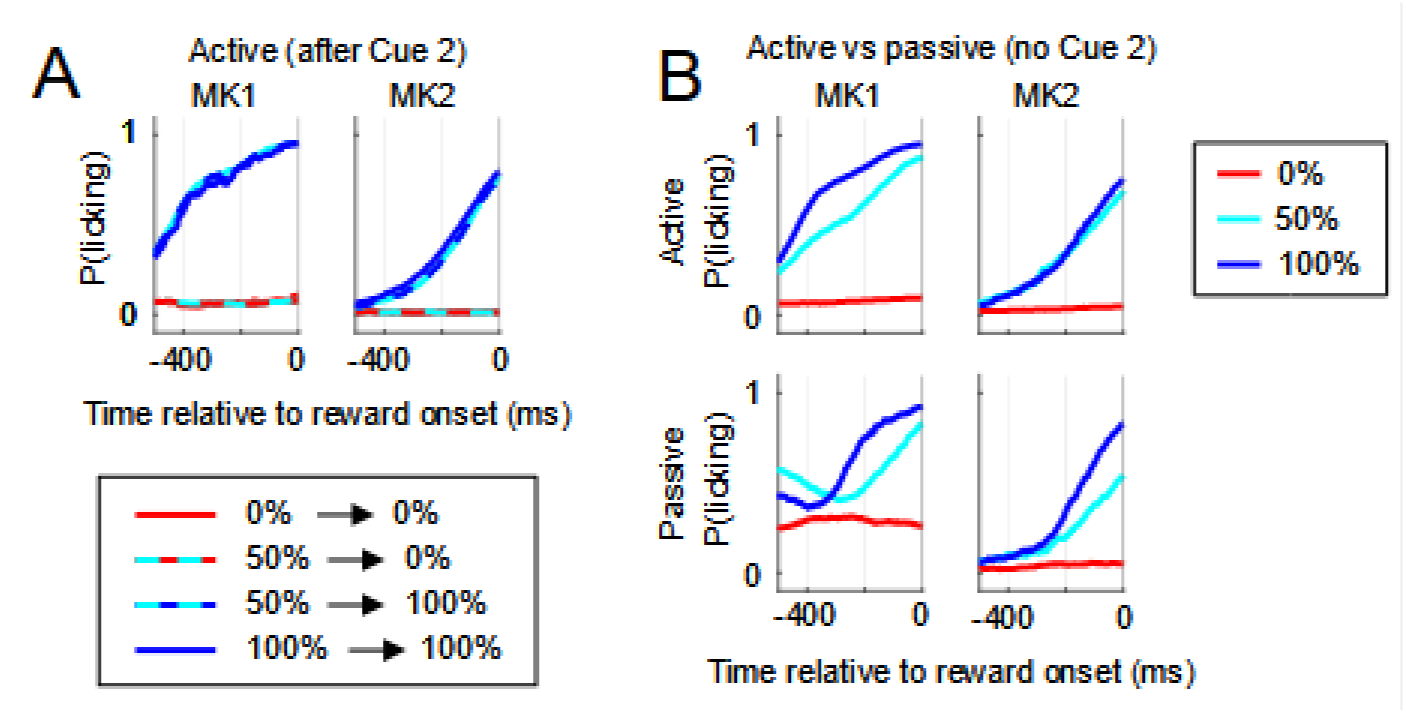
Anticipatory licking behavior shows that the monkeys were familiar with and attended to the cued reward probabilities. A. Probability of licking before reward onset after the monkeys revealed Cue 2. Both monkeys licked if Cue 2 signaled a reward but not if it signaled a lack of reward. **B.** Probability of licking in the absence of Cue 2 scales with the probability signaled by Cue 1 in both monkeys. The top row shows no-reveal trials on the active task (1-way ANOVA, p < 10^-43^, n = 37 in MK1, p < 10^-11^, n = 31 in MK2). The bottom row shows trials on the passive task when monkeys only viewed Cue 1 while fixating (1-way ANOVA, p < 10^-11^, n = 21 in MK1, p < 10^-5^, n = 20 in MK2).

**Supplementary Figure 2.**
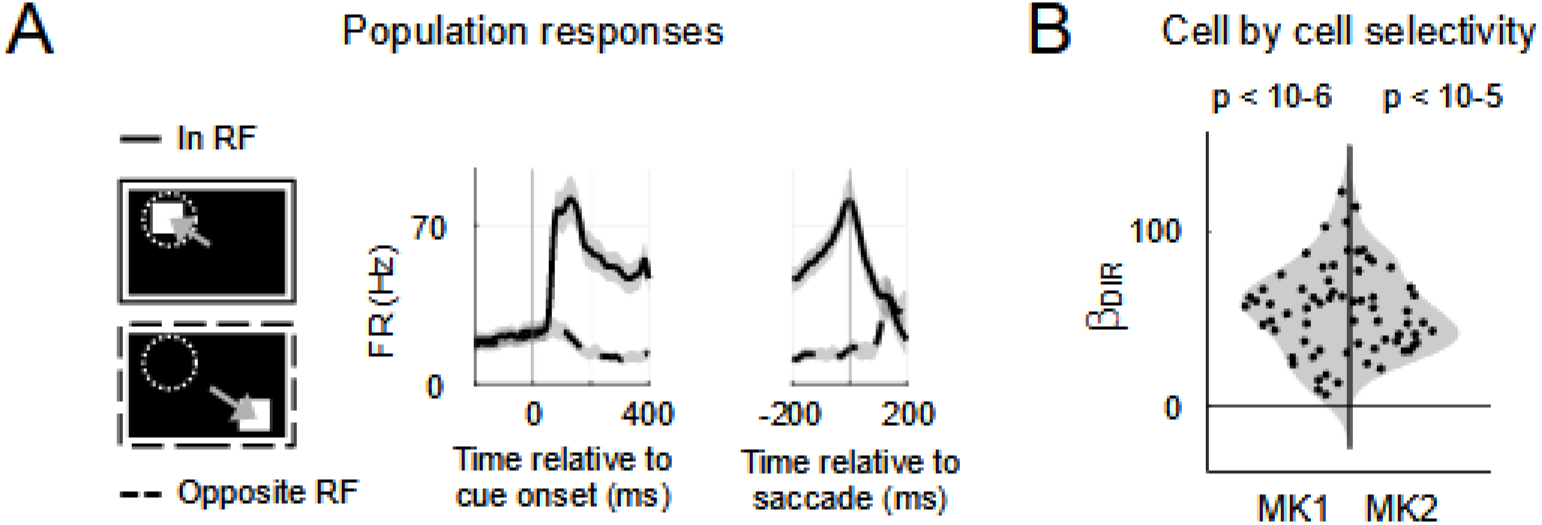
The recorded cells were spatially selective. A. PSTHs showing firing rates in the memory-guided saccade task, for target location/saccade goals inside the RF (solid) and opposite the RF (dashed; mean and 2 SEM, n = 68 cells). **B** Regression coefficients measuring saccade selectivity (FR ~ 1 + DIR, where DIR = 1 (0) if the saccade goal was inside (opposite) RF) in the 100 ms before saccade. Each point is one cell. The distributions (shading) were well above zero (signed-rank test against 0, MK1: p < 10^-6^, n = 37; MK2: p < 10^-5^, n = 31), and coefficients were individually significant in all but one cell in MK1 (which only showed spatially tuned visual but not pre-saccadic activity).

**Supplementary Figure 3.**
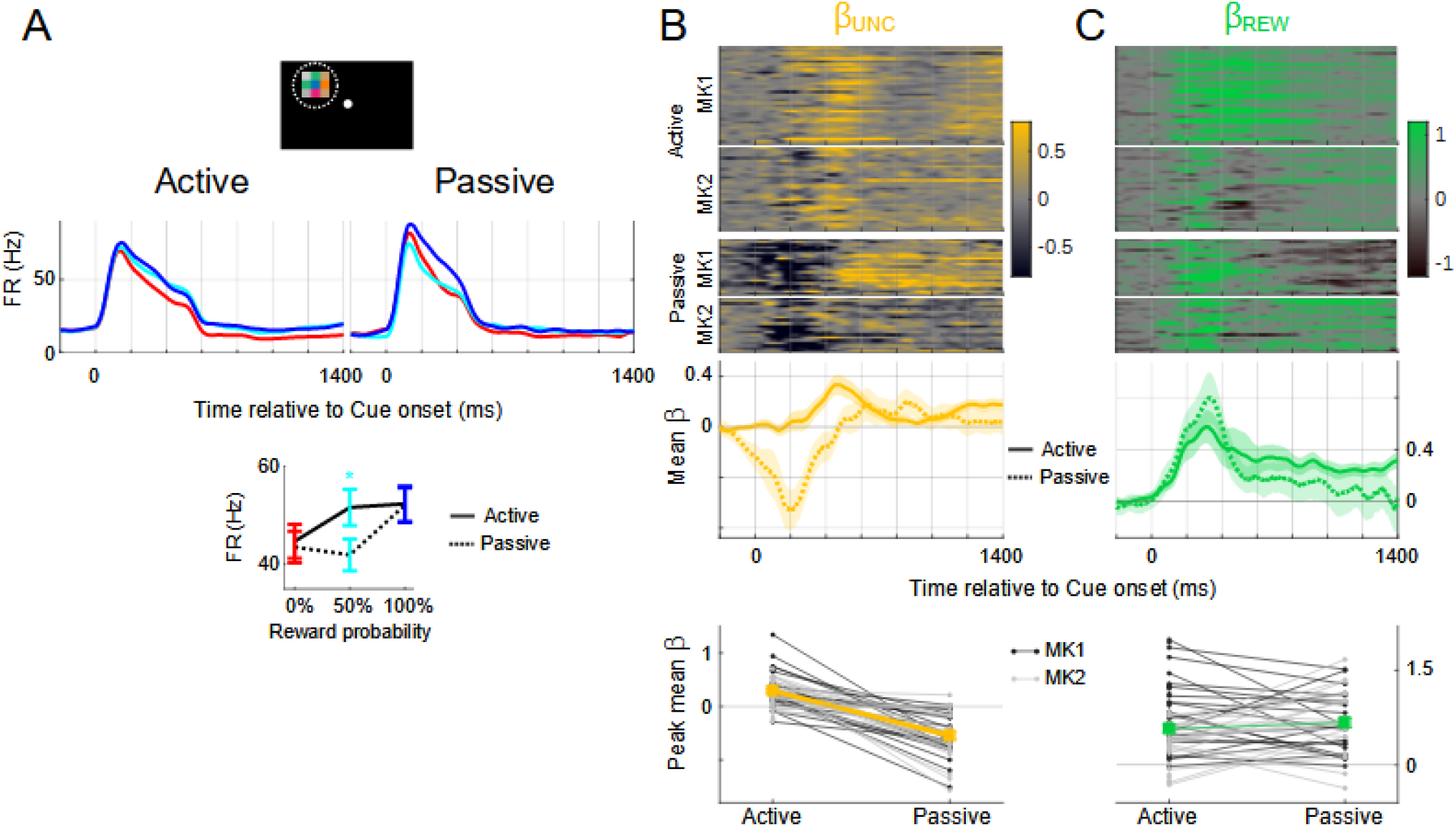
Context differentially affects uncertainty and reward modulations. A. PSTHs of responses to Cue 1 in the Active task (duplicated from **Fig. 2A**) and a “Passive” task in which the monkeys did not obtain visual information (*Methods*). Both tasks started with a period of central fixation during which Cue 1 stimuli indicated 0%, 50% or 100% reward probability. Anticipatory licking in the Passive condition scaled reliably with the probability (**Fig. S1C**), confirming that the monkeys attended to the Cue 1 information. When Cue 1 signaled 0% or 100% probability, the neurons had equivalent responses in the active versus passive condition. However, if the cue signaled 50% probability, firing rates were significantly lower in the passive condition (bottom, p < 10^-5^, n = 41 cells). Model based analysis confirmed that task context differentially affected uncertainty and reward modulations. **B.** In the Active condition, the peak β_UNC_ coefficient was significantly positive (200 ms window centered on average peak time (MK1, 0.34±0.05, n = 37; MK2, 0.24±0.04, n=31; both p < 10^-4^ relative to 0), while in the Passive condition, it was significantly negative (MK1, -0.54±0.09, n = 21, MK2, -0.55±0.11, n = 20, both p < 0.001 relative to 0). As shown by the colormaps and bottom plot, the difference was highly consistent in each of the cells tested in both tasks (paired signed-rank test, MK1: p < 10^-4^, n = 21; MK2, p < 0.001, n = 20 cells). **C.** In contrast, β_REW_ coefficients were positive and of similar or slightly higher magnitudes in the Passive relative to the Active task. A 2-way ANOVA revealed a significant interaction between signal and task (MK1: p < 0.01, MK2, p < 10^-8^; both p 10^-8^), confirming that task context distinctly affected the neuronal sensitivity to reward and uncertainty.

**Supplementary Figure 4.**
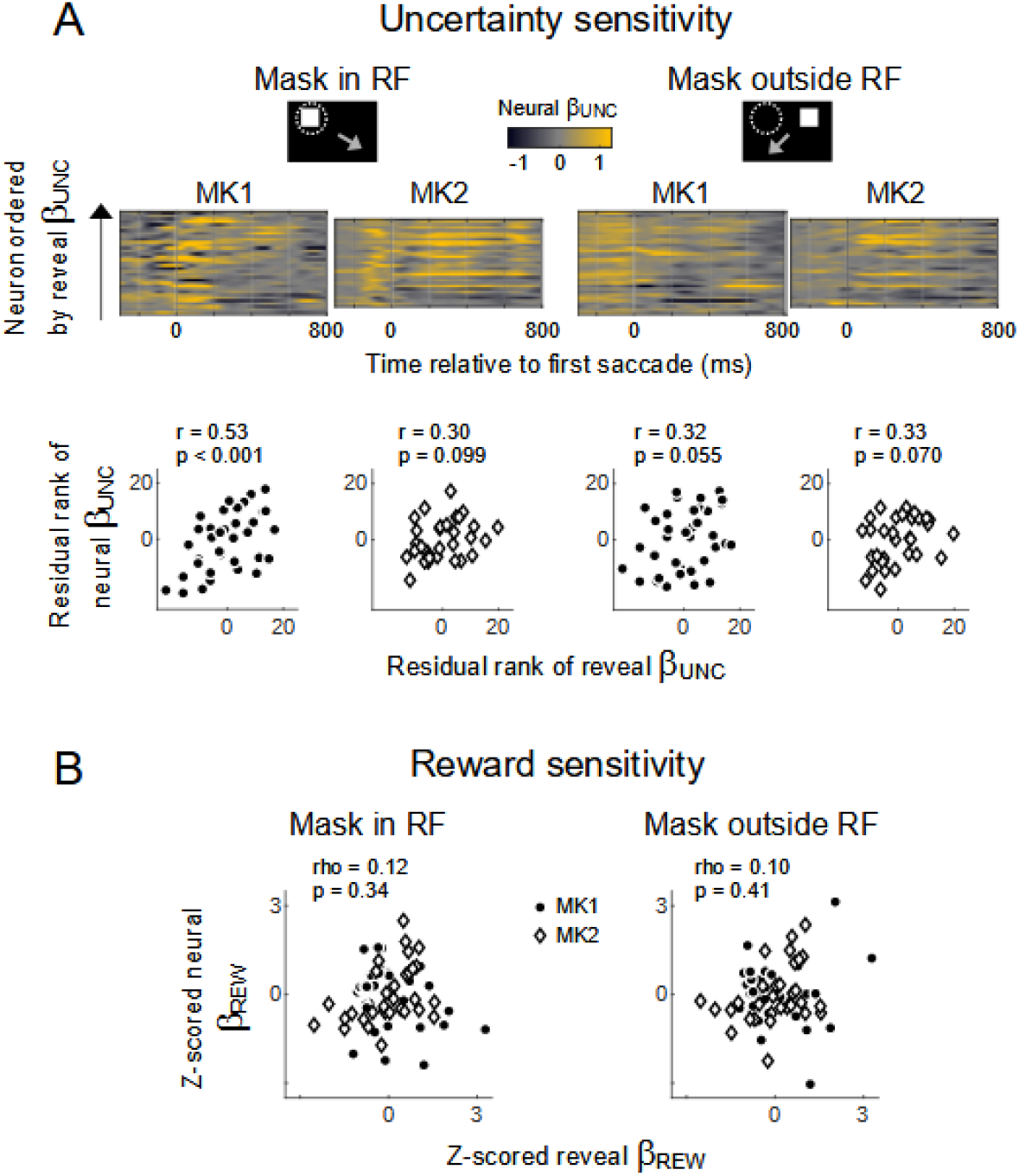
Neural-behavioral correlations remain consistent in all trials. **A.** Correlations between neural and behavioral β_UNC_ were generally replicated when we included reveal trials. Note that, when including reveal trials, post-saccadic neural responses encode the revealed information (**Fig. 5**), adding noise that accounts for the marginally significant effects in MK2. **B.** Correlations for neural and reveal β_REW_ coefficients are again not significant.

## References

1 Gottlieb, J. & Oudeyer, P. Y. Toward a neuroscience of active sampling and curiosity. Nat Rev Neurosci. 19, 758–770 (2018).

2 Bromberg-Martin, E. S. & Sharot, T. The Value of Beliefs. Neuron 106, 561–565, doi:10.1016/j.neuron.2020.05.001 (2020).

3 Daddaoua, N., Lopes, M. & Gottlieb, J. Intrinsically motivated oculomotor exploration guided by uncertainty reduction and conditioned reinforcement in non-human primates. Sci Rep 6, doi:doi: 10.1038/srep20202. (2016).

4 Kobayashi, K., Ravaioli, S., Baranès, A., Woodford, M. & Gottlieb, J. Diverse motives for human curiosity. Nat Hum Behav. 3, 587–595, doi:10.1038/s41562-019-0589-3 (2019).

5 Charpentier, C. J., Bromberg-Martin, E. S. & T., S. Valuation of knowledge and ignorance in mesolimbic reward circuitry. Proc Natl Acad Sci U S A. 115, E7255–E7264 (2018).

6 Dayan, P., Kakade, S. & Montague, P. R. Learning and selective attention. Nat Neurosci 3 Suppl, 1218–1223, doi:10.1038/81504 (2000).

7 Yang, S. C., Lengyel, M. & Wolpert, D. M. Active sensing in the categorization of visual patterns. eLife, doi:pii: e12215. doi: 10.7554/eLife.12215. (2016).

8 Squire, R., Noudoost, B., Schafer, R. & Moore, T. Prefrontal contributions to visual selective attention. Annu Rev Neurosci 8, 451–466, doi:doi: 10.1146/annurev-neuro-062111-150439. (2013).

9 Bisley, J. W. & Goldberg, M. E. Attention, intention, and priority in the parietal lobe. Annual Review of Neuroscience 33, 1–21 (2010).

10 Horan, M., Daddaoua, N. & Gottlieb, J. Parietal neurons encode information sampling based on decision uncertainty. Nat Neurosci 22, 1327–1335, doi:10.1038/s41593-019-0440-1 (2019).

11 Ebitz, R. B., Albarran, E. & Moore, T. Exploration disrupts choice-predictive signals and alters dynamics in prefrontal cortex. Neuron 97, 450–461 (2018).

12 Silvetti, M., Horan, M., LaSaponara, S. & Gottlieb, J. A reinforcement meta learning account of information demand and free energy. in preparation (2021).

13 Monosov, I. E. & Hikosaka, O. Selective and graded coding of reward uncertainty by neurons in the primate anterodorsal septal region. Nat Neurosci 16, 756–762, doi:10.1038/nn.3398 (2013).

14 Ledbetter, N. M., Chen, C. D. & Monosov, I. E. Multiple Mechanisms for Processing Reward Uncertainty in the Primate Basal Forebrain. J Neurosci 36, 7852–7864, doi:10.1523/JNEUROSCI.1123-16.2016 (2016).

15 Schultz, W. et al. Explicit neural signals reflecting reward uncertainty. Philos Trans R Soc Lond B BiolSci 363, 3801–3811, doi:J3K6217356280157 [pii]10.1098/rstb.2008.0152 (2008).

16 Jezzini, A., Bromberg-Martin, E. S., Trambaiolli, L. R., Haber, S. N. & Monosov, I. E. A prefrontal network integrates preferences for advance information about uncertain rewards and punishments. Neuron in press (2021).

17 White, J. K. et al. A neural network for information seeking. Nat Commun 10, 5168, doi:10.1038/s41467-019-13135-z (2019).

18 Louie, K., Grattan, L. E. & Glimcher, P. W. Reward value-based gain control: divisive normalization in parietal cortex. J Neurosci 31, 10627–10639, doi:31/29/10627 [pii]10.1523/JNEUROSCI.1237-11.2011 (2011).

19 Peck, C. J., Jangraw, D. C., Suzuki, M., Efem, R. & Gottlieb, J. Reward modulates attention independently of action value in posterior parietal cortex. J Neurosci 29, 11182–11191, doi:29/36/11182 [pii]10.1523/JNEUROSCI.1929-09.2009 (2009).

20 Ipata, A. E., Gee, A. L., Gottlieb, J., Bisley, J. W. & Goldberg, M. E. LIP responses to a popout stimulus are reduced if it is overtly ignored. Nat Neurosci 9, 1071–1076 (2006).

21 Reynolds, J. H. & Heeger, D. J. The normalization model of attention. Neuron 61, 168–185, doi:S0896-6273(09)00003-8 [pii]10.1016/j.neuron.2009.01.002 (2009).

22 Sugrue, L. P., Corrado, G. S. & Newsome, W. T. Choosing the greater of two goods: neural currencies for valuation and decision making. Nat Rev Neurosci 6, 363–375 (2005).

23 Leong, Y., Radulescu, A., Daniel, R., DeWoskin, V. & Niv, Y. Dynamic Interaction between Reinforcement Learning and Attention in Multidimensional Environments. Neuron 93, 451–463 (2017).

24 Noudoost, B. & Moore, T. Control of visual cortical signals by prefrontal dopamine. Nature,doi:nature09995 [pii]10.1038/nature09995 (2011).

25 Yu, A. J. & Dayan, P. Uncertainty, neuromodulation, and attention. Neuron 46, 681–692, doi:S0896-6273(05)00362-4 [pii]10.1016/j.neuron.2005.04.026 (2005).

26 Thiele, A. & Bellgrove, M. A. Neuromodulation of Attention. Neuron 97, 769–785, doi:10.1016/j.neuron.2018.01.008 (2018).

27 Gottlieb, J. & Snyder, L. H. Spatial and non-spatial functions of the parietal cortex. Curr Opin Neurobiol 20, 731–740, doi:S0959-4388(10)00185-6 [pii]10.1016/j.conb.2010.09.015 (2010).

28 Foley, N. C., Kelley, S. P., Mhatre, H., Lopes, M. & Gottlieb, J. Parietal neurons encode expected gains in instrumental information. Proceedings of the National Academy of Science 114, E3315–E3323 (2017).

29 Purcell, B. A., Schall, J. D., Logan, G. D. & Palmeri, T. J. From salience to saccades: multiple-alternative gated stochastic accumulator model of visual search. Journal of Neuroscience 32, 3433–3446 (2012).

30 Pouget, P. et al. Neural basis of adaptive response time adjustment during saccade countermanding. J Neurosci 31, 12604–12612, doi:31/35/12604 [pii]10.1523/JNEUROSCI.1868-11.2011 (2011).

31 Oristaglio, J., Schneider, D. M., Balan, P. F. & Gottlieb, J. Integration of visuospatial and effector information during symbolically cued limb movements in monkey lateral intraparietal area. J Neurosci 26, 8310–8319 (2006).

32 Foley, N. C., Cohanpour, M., Semework, M., Sheth, S. A. & Gottlieb, J. Population coding of reward prediction errors through opponent organization in the fronto parietal network. doi: https://doi.org/10.1101/769869(2020).

